# Construction of Feed Forward MultiLayer Perceptron Model For Genetic Dataset in Leishmaniasis Using Cognitive Computing

**DOI:** 10.1101/411363

**Authors:** Sundar Mahalingam, Ritika Kabra, Shailza Singh

## Abstract

Leishmaniasis is an endemic parasitic disease, predominantly found in the poor locality of Africa, Asia and Latin America. It is associated with malnutrition, weak immune system of people and their housing locality. At present, it is diagnosed by microscopic identification, molecular and biochemical characterisation or serum analysis for parasitic compounds. In this study, we present a new approach for diagnosing Leishmaniasis using cognitive computing. The Genetic datasets of leishmaniasis are collected from Gene Expression Omnibus database and it’s then processed. The algorithm for training and developing a model, based on the data is prepared and coded using python. The algorithm and their corresponding datasets are integrated using TensorFlow dataframe. A feed forward Artificial Neural Network trained model with multi-layer perceptron is developed as a diagnosing model for Leishmaniasis, using genetic dataset. It is developed using recurrent neural network. The cognitive model of the trained network is interpreted using the maps and mathematical formula of the influencing parameters. The credit of the system is measured using the accuracy, loss and error of the system. This integrated system of the leishmaniasis genetic dataset and neural network proved to be the good choice for diagnosis with higher accuracy and lower error. Through this approach, all records of the data are effectively incorporated into the system. The experimental results of feed forward multilayer perceptron model after normalization; mean square error (219.84), loss function (1.94) and accuracy (85.71%) of the model, shows good fit of model with the process and it could possibly serve as a better solution for diagnosing Leishmaniasis in future, using genetic datasets.

The code is available in Github repository:

https://github.com/shailzasingh/Machine-Learning-code-for-analyzing-genetic-dataset-in-Leishmaniasis

## INTRODUCTION

Leishmaniasis is the tropical protozoan parasitic skin disease. It is transmitted through the bite of sand flies, *Phlebetomine* sp. ^1^. It is found widely in the tropical and sub-tropical regions of the World ^2^. About 2 million new cases were emerging every year and they also lead to fatal in some cases. It mainly affects the poor locality of Asia, Africa and America. This is due to the population displacement, lack of sufficient nutrient and immunity in human beings and also their local pollutions. 97 out of 200 countries are endemic to leishmaniasis. Leishmania is a protozoal parasite responsible for Leishmaniasis. There are about 20 *Leishmania* sp., causing major leishmanial infection in human beings ^3^. There are three common types of leishmaniasis; Cutaneous leishmaniasis, Muco-cutaneous leishmaniasis and Visceral leishmaniasis (Kala-azar (or) Black fever) ^4^. Leishmanial parasites are capable of modulating the human immune system for their survival. They also modify the self-healing cutaneous lesions to fatal conditions ^5,6^. And some species of leishmaniasis are becoming virulent and drug resistant. These features of leishmaniasis made the researchers to focus on the comprehensive identification and analysis of its metabolic pathway and genome^1^.

Leishmaniasis is diagnosed by 4 different methods approved by US Centre for Disease Control and prevention. They are slide specimen, culture medium, Polymerase Chain Reaction (PCR) and serological testing. The biopsy specimens, dermal pus (or) scraps and impression smears are tested as slide specimens. The specialized culture medium is also used for the diagnosis of Leishmaniasis. The species identification of the leishmaniasis is carried out using PCR, with the complementary probes. The serum samples were also tested using the rK39 rapid test kit for diagnosing Leishmaniasis ^7^.

The medical data were significantly managed and secured using computer and networking in the past decades. The increase in the number of patient’s data lead to the new technology and technique called data mining, helpful in using a processed information. Its evolution lead it to the new field named health informatics, which have been used in the health sector for diagnosing the disease states and maintaining its data. These extended applications of data mining and cognitive computing have paved a way for the physician to diagnose the disease easily and fast, which have reduced the critical side effects on major diseases. They are also cost effective and more accurate ^8^. The diagnosis of disease using Artificial Neural Network (ANN) of feed forward back propagation networks have yielded a positive model for prediction. The working of biological neuron is mimicked using ANN, which process the distributed data in a parallel fashion. It stores and passes the acquired knowledge of data from one neuron to the other for responding to the given situation (or) conditions. ANN is used in normalizing the variance of data, which forms a linear robust model of data. ANN is better than the conventional multiple regression model, which cannot associate the non-linearity of the biological data. ANN can associate the multidimensional non-linearized data into a robust model using normalization factors ^9^.

Recurrent Neural Network (RNN) is a fully connected network, similar to feed forward neural network. But, the activation of each layer for RNN is dependent on the output of the previous layer. The temporary output memory of the RNN in passing the activation function leads it to be a good fit model for the prediction of robust data. It can also be activated by the sigmoidal activation function, used in normalizing the deviations in data. The input feed data of the RNN is three dimensional, stating the sample features, sample size and time series of length ^10^.

Early diagnosis of leishmaniasis would reduce the mortality rate and control the infectious stages of the disease. Hence, it needs the fast diagnosing model for the control of disease. This work emphasizes on the importance of Datamining and Cognitive computing in the diagnosis of Leishmaniasis. We have proposed the medical diagnostic scheme for diagnosing leishmaniasis, using genetic datasets of leishmaniasis. The genetic dataset was processed using RNN. RNN is used to recognize the variance in the genetic dataset of leishmaniasis and to form a robust dataset defining model using normalization process. This trained model will be applied for the diagnosis of leishmaniasis, using genetic dataset.

## Methodology

The varying factors (features) are essential to study the pattern of the given dataset accurately. The Genetic datasets of Leishmaniasis are collected from the public domain. All the records of the domains are experimentally verified. The aim of this work is to predict the valuable robust dataset defining model of the given genetic datasets. The variabilities of the data were used as features and attributes of algorithm, used in constructing a model for leishmanial diagnosis.

The flow of the work with the leishmaniasis genetic dataset to produce the robust model of feed forward multilayer perceptron, useful in diagnosing leishmaniasis, is represented in Figure 1. The raw genetic datasets are collected from the Gene Express Omnibus database. It is then processed to form a single dataset compiling all samples data with label encoded for each sample. It is processed through a feed forward multilayer perceptron model, using TensorFlow dataframe with the developed python code. The model is trained, checked for its accuracy, loss function and mean square error of the model at each iteration.

**Figure 1:**
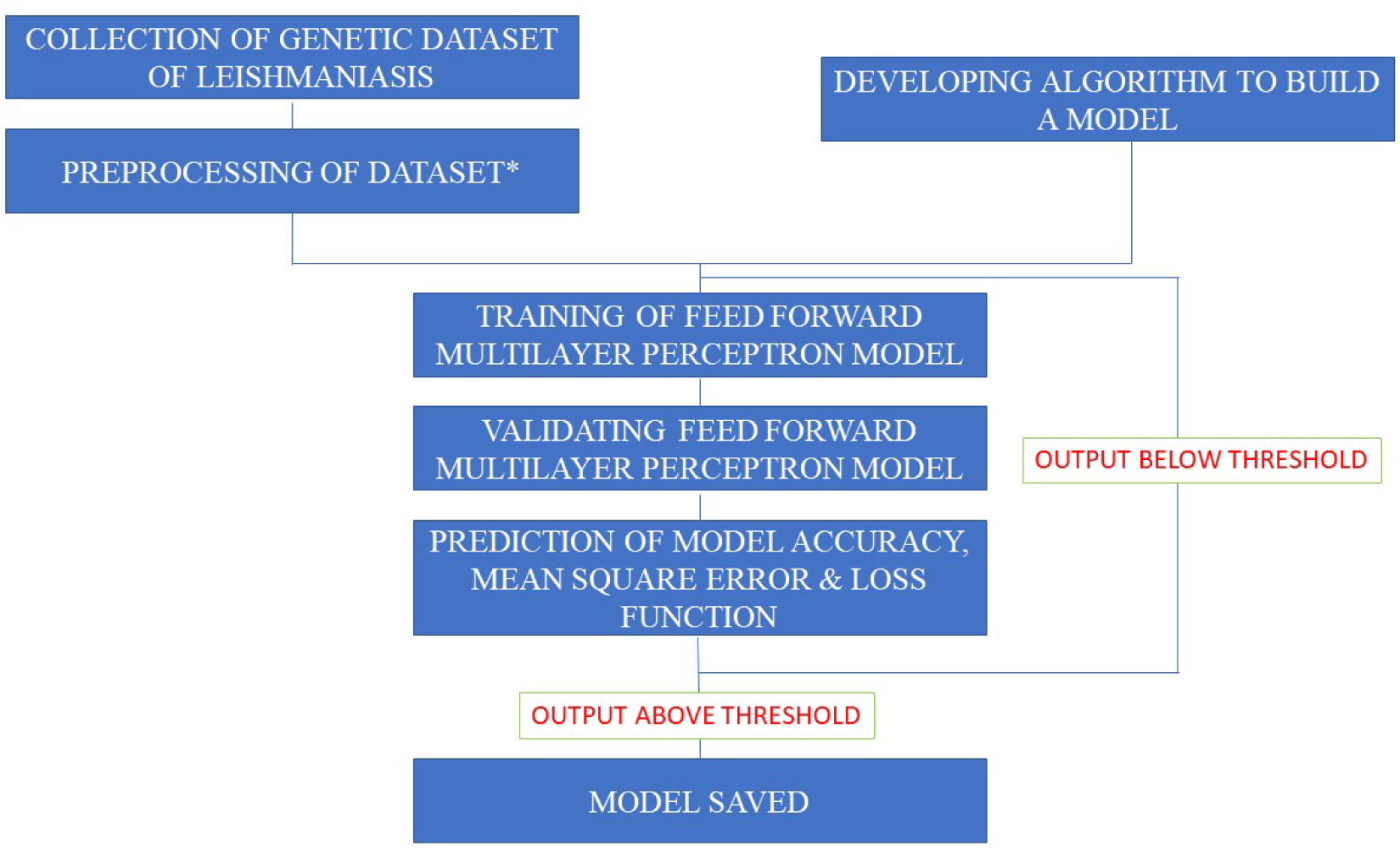
Schematic representation of the workflow designed.

*-The dataset with the same size of the features should be selected and processed

## Dataset

Gene Express Omnibus (GEO) database is one of the public data repositories of functional genomic data. It comprises of the array and sequence-based experimental gene expression data of all diseases. The 33 genetic datasets for total RNA and double stranded cDNA concentration of the leishmaniasis in human beings with same number of variables (45033 rows), done at same gene expression array analyser are collected. It comprises of 5 healthy person datasets and 28 infected human datasets. We labelled each dataset with the specific label code and compiled in a large single dataset in comma separated values file (.CSV) format. We used 80% of datasets as training data and the remaining 20% as the test data for the creation of the model.

## TensorFlow

The multidimensional array of the data is said to be tensor. The graphical and mathematical operations over the collected tensors are carried out through TensorFlow. It is an open source dataframe library. It comprises the libraries for various mathematical computation and graphical interpretations. The flow of numerical data over the mathematical operations are studied using the graphical representations. The large scale heterogenous distributed data of array are easily processed by the mathematical operations, which are coded as nodes in a processing algorithm. The major use of TensorFlow is to handle the big data of the dataset into a machine learning algorithm using suitable coding language ^11^. We have used TensorFlow 1.8.0 along with the Python 3.6 coding platform in this paper.

## Recurrent Neural Network

A feed forward ANN with multilayer perceptron is used as the bio-mimic neural network in this study to develop a model for diagnosing leishmaniasis, using genetic dataset. Every nodes of the ANN are used as the neurons of the human brain. The weights and biases of every node in the first layer is given with the input data. Then the output of every layer is passed as an input for the successive layers. We have used 4 hidden layers in this work. The multilayer perceptron involved in this process is used to study the non-linear functions of the given genetic data. Back propagation of data is also done using the code to increase the accuracy of prediction using the model, named as supervised learning of data. It compares the labelled data and the output data for its accuracy prediction. The rise or fall in the value is adjusted with the back-propagation process of ANN, until it reaches the threshold value of the process. This gradient descent optimization with the back-propagation process minimize the error. It is also efficient in finding the highly reproducible pattern of the given datasets ^11^.

## Neural Network Parameters

The various key terminology used in neural networks are listed below^8,12,13^.

- **Features**: The number of varying parameters and variables are stated as features (or) patterns (or) attributes.
- **Label**: The identity code of each sample to state its class
- **Weight**: The strength of the input variable denoted by numerical, involved in activation of neurons. It is adaptive, stating the connection between the two layers.
- **Biases**: The hidden matrix deciphered before all layers added up to the output for increasing the model efficacy
- **Input layer**: The layer with all the input variables for processing a model
- **Output layer**: The layer compiling all output and its corresponding interpretations
- **Hidden layer**: The layer involved in processing the input data to output, it lies between the input and output layer
- **True positive**: The leishmaniasis dataset correctly classified as leishmaniasis
- **True negative**: The dataset of normal healthy persons identified as normal
- **False positive**: The normal person dataset identified as leishmaniasis
- **False negative**: The leishmaniasis dataset identified as normal dataset

## Feed Forward Multilayer Perceptron Model

The model is created with the dataset which contains 33 samples with classes; leishmaniasis infected and normal healthy person genetic dataset of total RNA and double stranded cDNA concentration. The dataset is split into training (80%) and testing (20%). The model is trained at the learning rate of 0.001. The training is repeated for 500 epochs. We processed the data through Multilayer perceptron having 4 hidden layers with 45 neurons in each layer. The accuracy, cost (or) Loss function and Mean square error of the model is studied for each iterations and plotted as a graph. The variations in the data are normalized using log2 normalizations. The change in the threshold value is optimized using the Gradient Descent optimizer. The method is followed with the proposed method of Alagha et al., 2018, with slight modification to the operating parameters and hidden layer ^8^.

1. **Weight**: The output of each neuron depends on the weight matrix. The output of each neuron can be given as equation 1

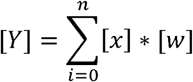

Where [Y] is the output matrix of corresponding layer, [x] is the input matrix; [w] is the weight matrix. Neural networks optimize the model learning through the iterative changing of the weight matrix values between the layers to attain the predefined threshold value. The iterative change in the weight value is carried out through the back propagation with respect to the defined dataset and threshold value ^12^
2. **Biases**: The extra matrix of the neural network lied before all input layers as pre-output layer, which is added to the matrix to get the final output of layer.
3. **Layer output**: The output of each layer is the sum of the multiplied matrix of input and weight matrix with the bias matrix

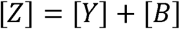 Where [Z] is the output of the layer and [B] is the bias matrix.
4. **Accuracy**: The accuracy of the model is the ratio of the total true predictions to the overall predictions ^8^. The accuracy of the model is given as

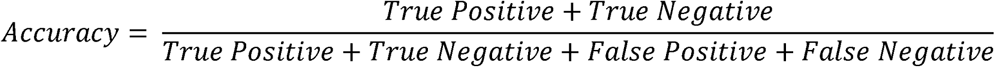
5. **Normalization**: The variations in the genetic dataset is normalized using log2 normalization function by leaving the least reproducing data ^8^. The normalization of the data process is mathematically denoted as

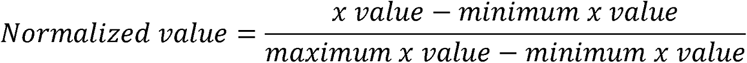
6. **Cost (or) Loss Function**: The loss function (or) cost of the model is calculated by the prediction of deviations in the mean squared error between the actual and predicted output value ^9^.
7. **Mean Square Error**: The mean of the total variations between the actual labelled output and the predicted output is measured as Mean Square Error ^14^.

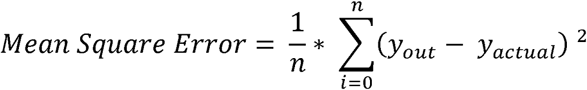
8. **Gradient Descent Optimizer**: Artificial neural network could not capture highly depending variables of genetic dataset, while back propagation. The gradient of the error function is decreased exponentially with number of epochs, using gradient descent optimizer and the highly reproduceable data are captured by long short term memory of the model ^15^.

## RESULTS AND DISCUSSION

### Feed Forward Multilayer Perceptron Model

The back propagation induced feed forward network with multilayer perceptron is activated using sigmoid function. The usage of back propagation in network have decreased the loss function to the prescribed threshold value by adjusting the weights of the neuron connections. The back propagation incorporated model with multilayer perceptron have added a good solution for the specific problem, diagnosis of Leishmaniasis using its genetic dataset. The lower learning rate of the process with the relative high accuracy of solution makes it as a good scheme of model for diagnosing leishmaniasis in human beings using the genetic dataset. The robust and stable model of the process have been developed using the back propagation, which is used to train a non-linear data into a trained model with mathematical notations. The accuracy, loss function and the mean square error of the corresponding epoch is tabulated in Table 1.

**Table 1:** Accuracy, Loss function and Mean Square Error of Training Epochs

| Epoch No. | Accuracy (%) | Loss Function | Mean Square Error |
| --- | --- | --- | --- |
| 0 | 0 | 9.91 | 245.96 |
| 25 | 14.29 | 7.46 | 229.82 |
| 50 | 28.57 | 5.37 | 219.81 |
| 75 | 42.86 | 3.86 | 214.99 |
| 100 | 42.86 | 3.17 | 213.62 |
| 125 | 42.86 | 2.86 | 213.57 |
| 150 | 57.14 | 2.67 | 213.98 |
| 175 | 57.14 | 2.53 | 214.58 |
| 200 | 71.43 | 2.43 | 215.25 |
| 225 | 71.43 | 2.36 | 215.94 |
| 250 | 85.71 | 2.29 | 216.60 |
| 275 | 85.71 | 2.24 | 217.20 |
| 300 | 85.71 | 2.19 | 217.74 |
| 325 | 85.71 | 2.15 | 218.21 |
| 350 | 85.71 | 2.12 | 218.60 |
| 375 | 85.71 | 2.08 | 218.94 |
| 400 | 85.71 | 2.05 | 219.21 |
| 425 | 85.71 | 2.02 | 219.41 |
| 450 | 85.71 | 1.99 | 219.60 |
| 475 | 85.71 | 1.96 | 219.74 |
| 500 | 85.71 | 1.94 | 219.84 |

These characters of optimization are explained by Jin et al., 2018. The gradient decrease in the Loss function and Mean square error with respect to their corresponding epoch number is due to the Gradient Descent optimizer ^16^. This have also increased the accuracy of the model in prediction of leishmaniasis. The model is developed with 45 neurons. It has 4 hidden layers fully interconnected to each other, classifying two different classes of leishmaniasis. The schematic representation of the working model is represented in Figure 2.

**Figure 2:**
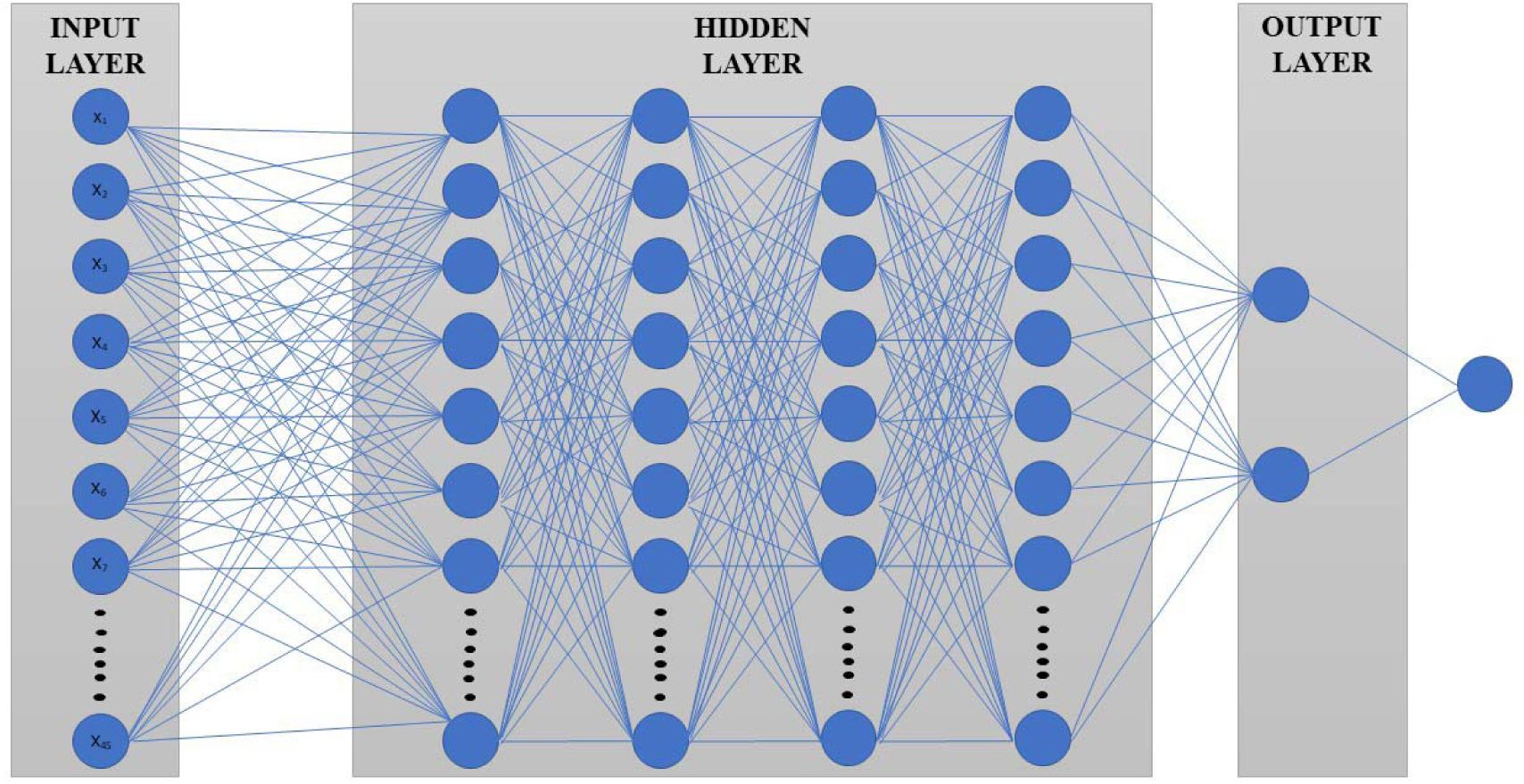
Schematic representation of the feed forward multilayer perceptron model

The 45033 features of the individual sample are loaded into the 45 neurons of the input layer. The datasets of 33 different human samples are processed into the input layer. The processed input is passed into the series of hidden layers for its data optimization, to select the maximum reproducing data of the given genetic dataset. It is then characterized into the two classes in the output layer. The value of the output class crossing above the threshold value is given as an output of the given test data. The random structure of hidden layer used in constructing the model increases the efficacy of the model, which is proved with increased accuracy of the model. The randomization of weight input with the hidden layers without affecting convergence would increase the fit of model with the process ^17^. The number of hidden layers required for the model plays a vital role in optimization of the model to act as a good efficient model. The usage of hidden matrix, bias, made the perfect stable model, without affecting its convergence ^18^. The increased accuracy of the model is due to the use of increased neurons in the hidden layer. The increased number of hidden layer, decreases the error of the model to the process ^19^. The input of each hidden layer is dependent on the output of the preceding neural layer. This dependency structure of the matrix made the model to learn the highest data volume of the genetic dataset and also it increases the cognitive effect of the model ^20^.

- **Accuracy:** The accuracy of the model is initially 0%. Then it started learning slowly with the increase in the number of iterations with epochs. They increase as a step by step impulse as through the learning and normalization of data occurring at each stages of the process. Then, the final accuracy of the data is attained to be 85.71%. showing it to be a good fit of model with the process. The variations in the accuracy through learning with their corresponding epoch (or) cycle is denoted in the Figure 3.

**Figure 3:**
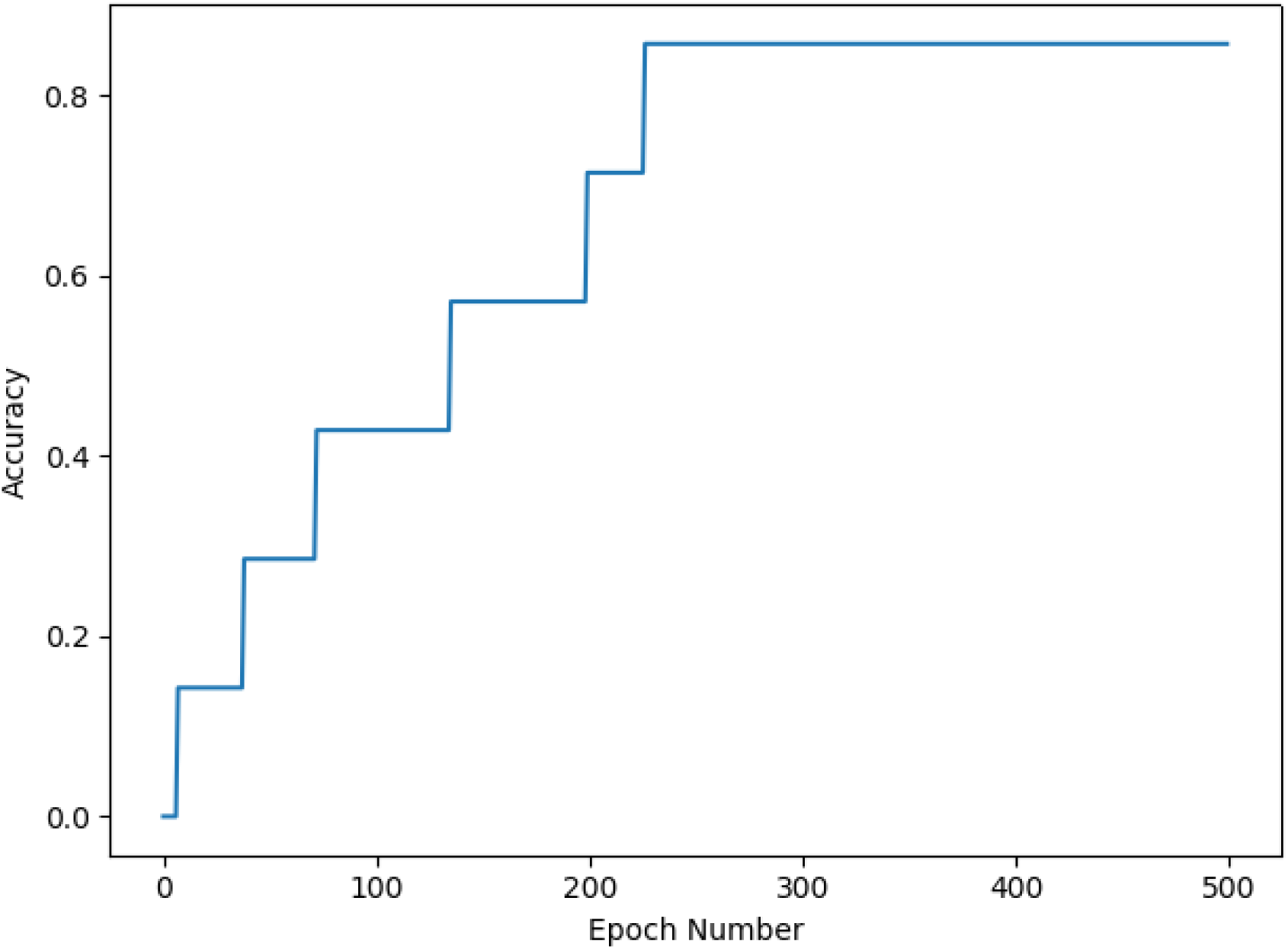
Effect of accuracy with their corresponding Epoch number The accuracy of 85.71% shows it to be a good model for predicting the more reliable experimental result for the given genetic dataset of leishmaniasis in human beings. The increase in the accuracy states the increase in the controlling of the parameters and features with the prediction ^21^. The increasing value of the accuracy denotes the decreasing state of the error in the model, with the increase in regression coefficient ^22^. The usage of back propagation has increased the learning of the model, which in turn increased the accuracy of the model. The usage of back propagation along the feed forward neural network have also proved to be the better option than using spike timing-dependent plasticity model for biological datasets. The lower classification accuracy and training difficulty of the deep networks is overcomed with this feed forward multilayer perceptron model with back propagation ^23^. The accuracy value above 80% is accepted. Since, it could clearly classify and identify the class with its special features with 70 % of total accuracy ^24^.
- **Loss Function:** The loss function (or) cost of the model is showing a significant decrease in the value with the continuous increase in the number of epochs. The loss function is found to be 9.91 at the start of the training process. Then the increase in the number of cycle (or) epoch, decreases the loss function (or) cost, with continuous learning through the datasets repeatedly. The further increase in the number of cycle used for training the model using training dataset have decreased the loss function (or) cost of the model. The variations of the Loss function with their corresponding epoch are given in Figure 4.

**Figure 4:**
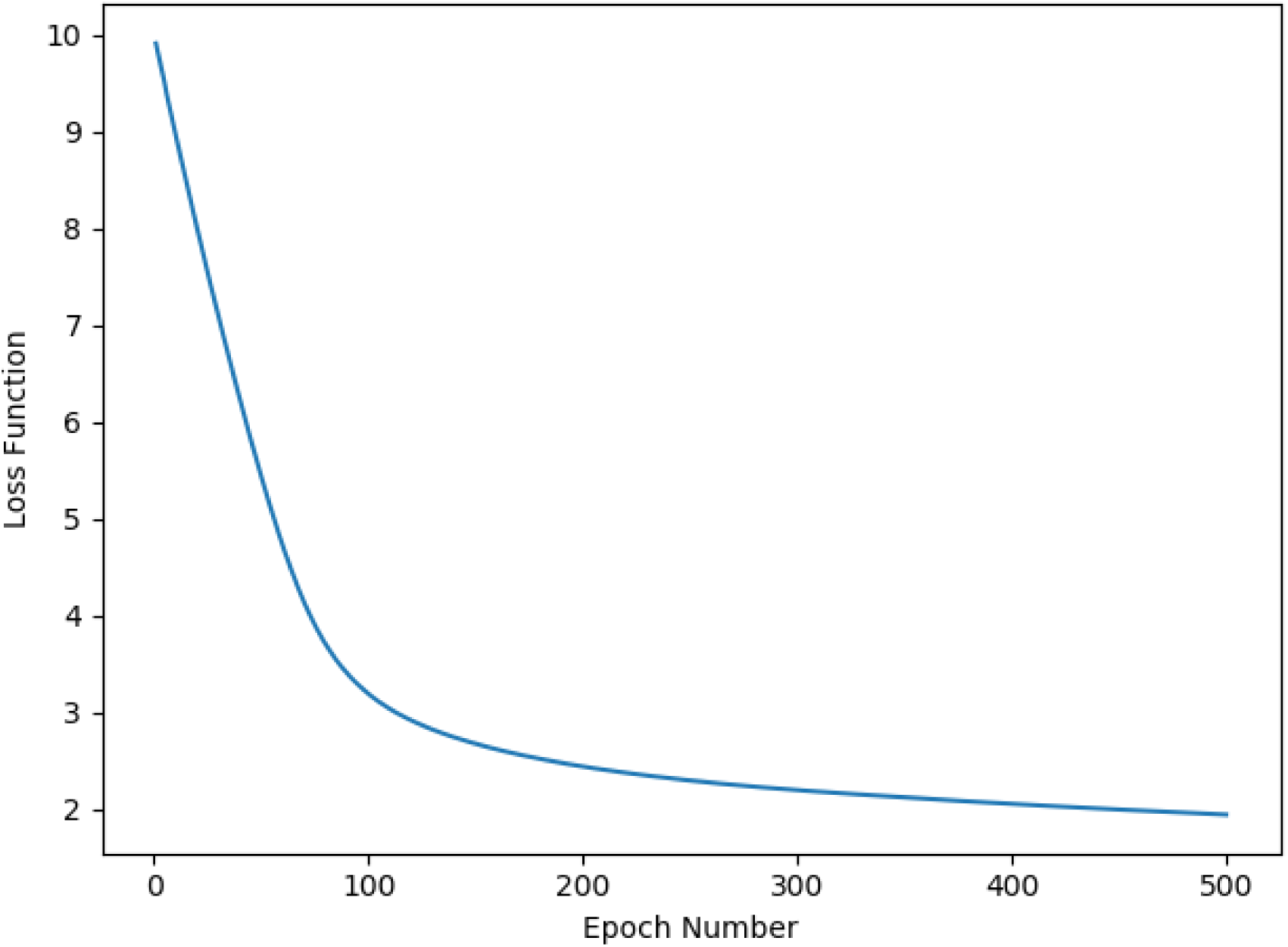
Effect of Loss Function with their corresponding Epoch number The intrusion of loss function into the model depicts the minor change of the dataset, with respect to its variations. It is also used in studying the impact of improvement. The improvement in the loss function shows the efficiency of the method in identifying the relatively indistinguishable data ^25^. The matching process of pairs is higher with the decrease in loss function. The decrease in loss function states the good fit of parameters with the developed model ^26^. The stability of the loss function shows the stability of the model with the given data. It also shows the improved functionality of parameters with reduced error ^27^. The decrease in the loss function shows the trained model with higher accuracy and it proves the idealistic property of the back propagation in learning. It is also used in proving the combination of parameter coefficients to be less with its improved values ^28^.
- **Mean Square Error:** The mean square error of the process is found to be 245.96 initially. There is a gradual decrease in the mean square error value till 100 epochs. Then there is a slight increase in the value from 100-300 epochs. Then it is standardized to a constant value of 219 after 300 epochs, which shows the normalized state of all data. The process of learning with the leishmaniasis genetic dataset is normalized to the yield of robust model, leaving out a least reproducing feature. The process of normalization is carried out in learning the dataset using the log 2 normalization methods. The variations in the mean square error with their corresponding epoch number are plotted in Figure 5.

**Figure 5:**
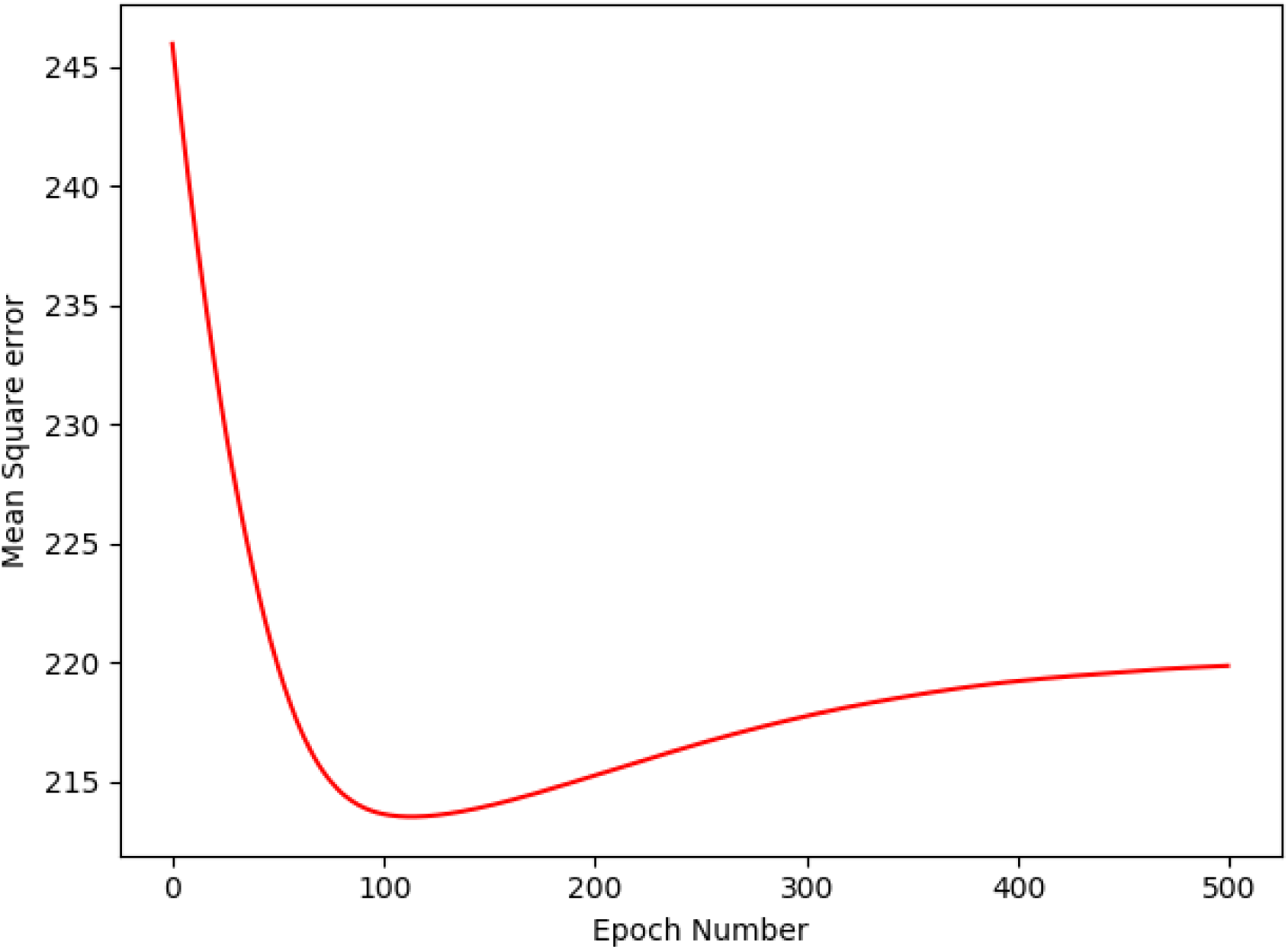
Effect of Mean Square Error with their corresponding Epoch number The smaller decrease of the mean square error is because of the high non-uniformity of the given dataset. The maintained level of deviations in the plot proves it to be normalized from higher deviations of data ^29^. The higher mean square error value is due to the lower samples of data ^30^. The decrease in the mean square error value also proves the working of feed-back gradient descent optimization mechanism of algorithm, through which the parameters are made to fit into the model correctly by improving the weight of the input data ^31^. It also proves the convergence of the data reliable to the robust model for the given dataset for the constant parameters ^32^ The mean square error is also used in studying the performance of the model with the neural network. The decrease in mean square error states the decrease in error between the network output and the target output, showing the good fit of model with the process ^33^.

### Conclusion

This study resulted with a new attempt in processing the genetic dataset of the human sample into the human disease diagnosis using cognitive computing. The hybridization of genetic dataset with its prediction model, created using algorithm, is a best solution for diagnosis of leishmaniasis in human beings in future using their genetic dataset. The use of feed forward multilayer perceptron model with back propagation have produced a good fit of model with the process of diagnosis, showing 85.71% of accuracy in prediction. By this we have brought a new solution of predicting disease in human beings using the genetic dataset with the integration of cognitive computing model. The classification of leishmaniasis in human beings using their genetic dataset of total RNA and c DNA concentration has resulted with the new model of prediction. The obtained result of accuracy, mean square error and loss function, proves the model to be good fit for the process of diagnosis in leishmaniasis using the genetic dataset. So, we suggest this model can be used for the diagnosis of leishmaniasis in human beings using their genetic dataset. This cognitive computing approach in diagnosis of Leishmaniasis in human beings, will also form a robust dataset emphasizing the maximum reproducing data, which could be a target set for the in-vitro researches regarding leishmaniasis in human beings. In future, the model would be trained with the large number of dataset, and thereby increasing the cognitive level of model in diagnosis and prediction of given dataset. It would also be made as an application for the leishmanial diagnosis in human beings using genetic dataset.

## Acknowledgement

We thank the Director, National Centre for Cell Science (NCCS), for supporting the Bioinformatics and High-Performance Computing Facility (BHPCF) at NCCS, Pune, India. Sundaramahalingam acknowledges the Indian Academy of Sciences, Bangalore, for offering a summer research fellowship. Also, acknowledges Secretary, Principal and Head of the Department, Department of Biotechnology, Kamaraj College of Engineering and technology, Madurai District for permitting him to take a fellowship during the course time. Ritika Kabra acknowledges her Senior Research Fellowship from Department of biotechnology, Ministry of Science and Technology, Government of India.

## Author’s Contribution

Sundar Mahalingam (SM) performed the building up of feed forward neural network model, statistically analysed the datasets, wrote the code and the manuscript. Ritika Kabra (RK) contributed in statistical analysis and building figures for the same. Shailza Singh (SS) conceptualized the whole idea, performed coding of the genetic datasets and wrote the manuscript.

